# Mitochondrial fission dysfunction alleviates heterokaryon incompatibility-triggered cell death in the industrial filamentous fungus *Aspergillus oryzae*

**DOI:** 10.1101/2021.12.10.472196

**Authors:** Chan Lu, Takuya Katayama, Noriko Mori, Ryota Saito, Kazuhiro Iwashita, Jun-ichi Maruyama

**Affiliations:** Department of Biotechnology, University of Tokyo, 1-1-1 Yayoi, Bunkyo-ku, Tokyo, 113-8657, Japan; Collaborative Research Institute for Innovative Microbiology, University of Tokyo, Bunkyo-ku, Tokyo, Japan; Division of Fundamental Research, National Research Institute of Brewing (NRIB), Hiroshima, Japan

**Keywords:** *Aspergillus oryzae*, heterokaryon incompatibility, strain phylogeny, mitochondrial fragmentation

## Abstract

In filamentous fungi, cell-to-cell recognition is a fundamental requirement for the formation, development, and maintenance of complex hyphal networks. Basically, self/compatible individuals within the fungal species are capable of fusing together, a step important for crossbreeding, which results in the formation of viable vegetative heterokaryons. Conversely, the fusion of incompatible individuals does not result in the formation of viable hyphal networks, but it often leads to growth inhibition or cell death. Even though a number of studies have been conducted to investigate such incompatibility, the understanding of the associated molecular mechanism is still limited, and this restricts the possibility of crossbreeding incompatible individuals. Therefore, in this study, the characteristics of compatibility/incompatibility in the industrial filamentous fungus, *Aspergillus oryzae*, were comprehensively investigated. Protoplast fusion and co-culture assays indicated the existence of a correlation between strain phylogeny and compatibility/incompatibility features. Time-course fluorescence observations were employed to investigate the types of incompatible responses that are induced at different cellular levels upon incompatible cell fusion, which eventually lead to cell death. Propidium iodide-indicated cell death, ROS accumulation, and mitochondrial fragmentation were identified as the major responses, with mitochondrial fragmentation showing the most significant subcellular change immediately after incompatible cell fusion. Furthermore, the deletions of mitochondrial fission-related genes *Aofis1* and *Aodnm1* in incompatible pairing alleviated cell death, indicating that mitochondrial fission is an important mechanism by which incompatibility-triggered cell death occurs. Therefore, this study provides new insights about heterokaryon incompatibility.

**IMPORTANCE:** For a long time, it was believed that as an asexual fungus, *A. oryzae* does not exhibit any sexual cycle. However, the fungus has two mating types, indicating the potential for sexual reproduction besides a known parasexual cycle. Therefore, given that viable heterokaryon formation following cell fusion is an important step required for genetic crossing, we explored the mechanism of incompatibility, which restricts the possibility of cell fusion in *A. oryzae*. Protoplast fusion and co-culture assays led to the identification of various vegetative compatible groups. Mitochondrial fragmentation was found to be the most significant incompatible cellular response that occurred in organelles during incompatible pairing, while the deletion of mitochondrial fission-related genes was identified as a strategy used to alleviate incompatibility-triggered cell death. Thus, this study revealed a novel mechanism by which mitochondrial fission regulates incompatible responses.

## INTRODUCTION

In filamentous fungi, cell fusion is vital for the development and maintenance of multicellular filament systems within vegetative individuals. Two fungal individuals capable of undergoing cell fusion form a heterokaryon, a multinucleate cell where genetically different nuclei co-exist (1). However, not all fungal individuals can safely fuse together. In particular, vegetatively incompatible individuals cannot form stable heterokaryons. They immediately reject each other at cell fusion, leading to programmed cell death or growth inhibition, which are termed vegetative incompatibility or heterokaryon incompatibility phenomena (2).

To measure fungal population diversity and appropriately characterize different individual isolates, vegetative compatible group (VCG) classification is usually applied, and over the past decades, vegetative incompatibility assays have been performed on numerous fungal species (3). For some model fungal species in particular, incompatibility is indicated by the formation of a barrage zone following the fusion of incompatible isolates (4, 5). Auxotrophic complementation assays in which only compatible complemented auxotrophic mutants can form prototrophic heterokaryons on minimal media, have been predominantly employed to discriminate compatible/incompatible relationships between the different isolates (3). For instance, VCG classification experiments based on complementation tests between nitrate-nonutilizing mutants have been extensively performed (6–9). Thus, it is often preferable to perform an auxotrophic complementation assay to determine VCGs given that it clearly demonstrates the ability of isolates to form heterokaryotic mycelia.

Additionally, it is widely believed that heterokaryon formation is regulated by the various molecular components that are associated with self/non-self discrimination (allorecognition) mechanisms, which primarily involve the interplay of protein products of the same (allelic) or different (non-allelic) loci of the *vic* (vegetative) and *het* (heterokaryon) loci (10). When any combination of incompatible alleles is co-expressed, the result is incompatibility-triggered cell death (10). The *het-S*/*het-s* system, which is a well-studied incompatibility system in *Podospora anserina*, acts as a prion causing the HET-S protein to become a pore-forming toxin that affects plasma membrane permeability and induce rapid cell death (11, 12). Reportedly, in *Neurospora crassa*, the deletion of the transcription factor gene *vib-1* suppresses the genetic difference-mediated incompatibility reaction at the *het* loci (13). In *P. anserina*, it has been observed that autophagy deficiency accelerates cell death, indicating that it plays a protective role against incompatibility-triggered cell death (14, 15). Although a number of genes involved in heterokaryon incompatibility in some filamentous fungal species have been characterized, the molecular basis of the cell death reaction is not well understood.

Incompatibility-triggered cell death is a rapid response that occurs within approximately 20 min in *N. crassa* incompatible cell fusion (16). During this process, plugged septal pores are a common feature that can spatially restrict the fused area of incompatible cells via increased compartmentation and intensive vacuolation (17–20). Transmission electron microscopic analysis has characterized the organelle degradation and disorganization associated with this process, and demonstrated the appearance of plasma membrane shrinkage/leakage (20). Additionally, reactive oxygen species (ROS) production, nuclear DNA degradation, and lipid droplet accumulation are typically observed during incompatible cell fusion (19, 21, 22). However, the correlation between incompatibility-induced physiological changes and the underlying mechanism is still unclear.

The filamentous fungus *Aspergillus oryzae* is widely used in the traditional Japanese brewing industry. Numerous strains of this fungus are selectively used for different purposes such as sake, soy sauce, and *miso* production, and the strains have been classified into 13 phylogenetic clades (23). Additionally, heterokaryon formation following cell fusion is an important step in genetic crossing associated with the industrial application of *A. oryzae*. However, the low cell fusion efficiency of the strains often makes the investigation of heterokaryon formation challenging. In our previous study, we established a biomolecular fluorescence complementation (BiFC) method to visualize fused cells and suggested the existence of heterokaryon incompatibility in *A. oryzae* (23). However, the overall strain phylogenetic preference and the dynamics of incompatible responses at cellular level remain elusive. In this study, we performed an auxotrophic complementation assay via protoplast fusion to investigate the incompatibility among 34 *A. oryzae* strains and clarified the phylogeny-preferred distribution. In addition, we optimized the co-culture conditions to significantly enhance fusion efficiency, which allowed us to successfully visualize the subcellular dynamics of the cell fusion process and the incompatible responses. Based on these findings, we first identified a correlation between mitochondrial fragmentation and incompatibility; subsequently, we demonstrated that incompatibility-triggered cell death can be alleviated by deleting mitochondrial fission-related genes.

## RESULTS

### Compatible group classification via auxotrophic complementation assay

In our previous study, we observed heterokaryon incompatibility in 14 *A. oryzae* strains using the BiFC technology, which involves visualizing fused cells in self- or non-self-pairings via cytoplasmic split proteins of enhanced green fluorescent protein (EGFP) (23). However, the study only focused on the incompatibility features by using the representative inter-clade strains. In addition, the BiFC method only represents transient fusion ability, and cell death resulting from incompatible fusion is, therefore, inconclusive.

To comprehensively investigate strain incompatibility, we selected 34 *A. oryzae* strains, which sufficiently represented all *A. oryzae* strain phylogenetic clades, including inter-clade and intra-clade strains (Fig. S1) (23). Auxotrophic complementation assays were performed using two uridine/uracil auxotrophic mutants of the genes *pyrF* (orotidine-5’-phosphate decarboxylase) and *pyrG* (cytidine triphosphate synthetase). We employed the CRISPR/Cas9 system to delete *pyrF* and *pyrG* genes in the 34 strains. If auxotrophically complemented heterokaryons grew on minimal medium under forced protoplast fusion between Δ*pyrG* and Δ*pyrF* strains, using polyethylene glycol (PEG), the two strains were classified into the same VCG. If the two strains did not produce heterokaryons, they were classified under different VCGs (Fig. S2A).

To analyze incompatibility features in inter-clade strains via the auxotrophic complementation assay, we selected 13 representative strains, which were derived from each of the phylogenetic clades (Fig. S1). Thirteen self-pairings clearly formed auxotrophic complemented mycelia on the minimal medium, which acted as a positive control (Figs. 1A and S2B). However, except for two pairs of phylogenetically close strains that showed heterokaryon growth, the other strain pairs showed no obvious growth (Figs. 1A and S2B). We subsequently examined intra-clade strains belonging to the same phylogenetic clades and observed the growth of mycelia in all the tested strain pairings (Figs. 1B and S2C). These incompatibility features corresponding to intra- and inter-clade strains revealed at least seven VCGs among the selected 34 *A. oryzae* strains (Fig. 1C).

**FIG 1.**
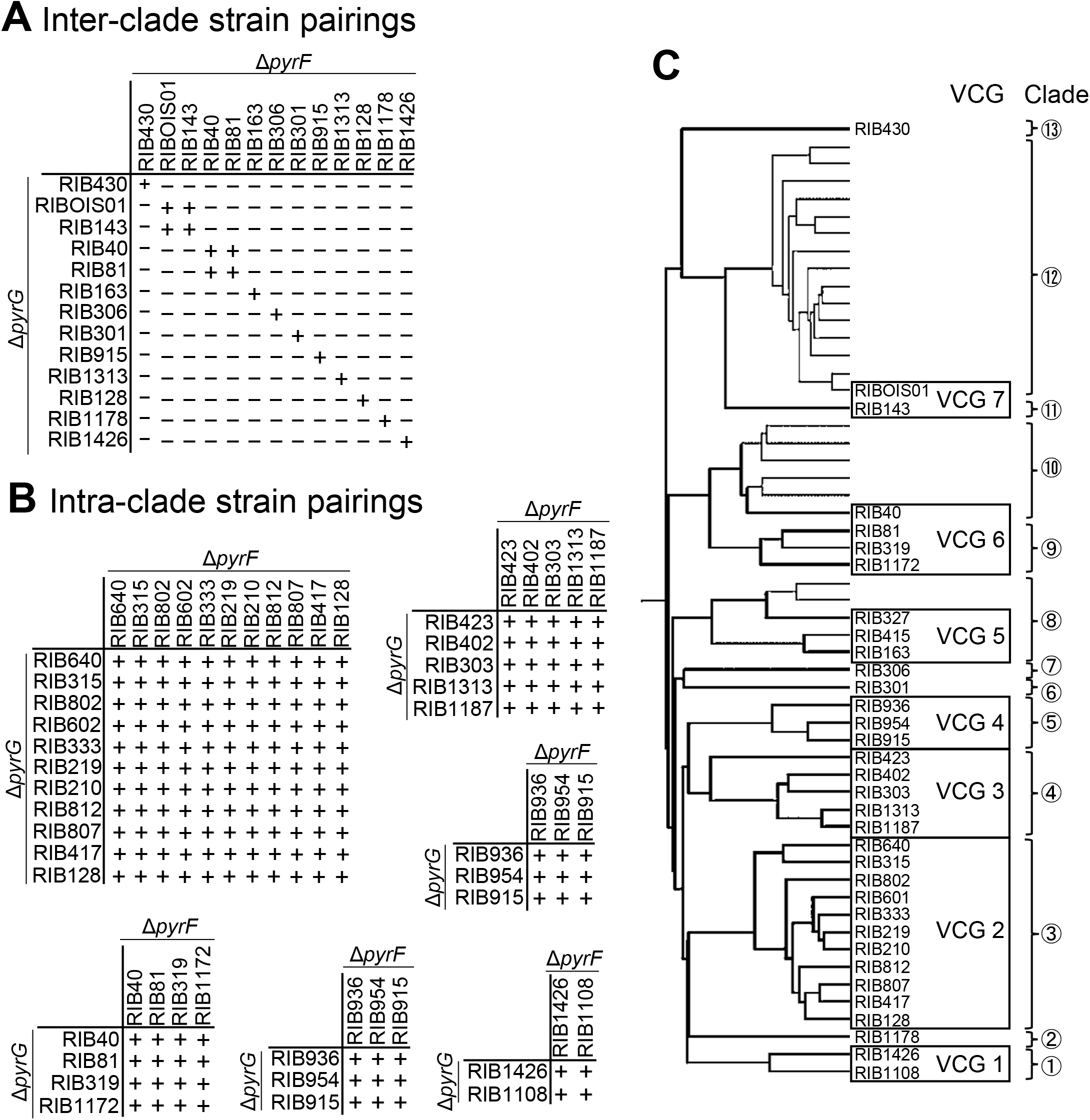
Correlation between heterokaryon incompatibility and strain phylogeny in *A. oryzae*. (A) Analysis of incompatibility between inter-clade strains. The 13 representative strains belong to 13 different clades. The detailed protocol and agar plate images are shown in Fig. S2A. (B) Analysis of incompatibility between intra-clade strains. The detailed protocol and plate images are shown in Fig S2A. (C) Vegetative compatible group (VCG) classification of compatible groups and strain phylogeny.

The VCG classifications compared to each strain phylogeny indicated that all the compatible strains belonged to the same phylogenetic clade or could be placed in close phylogenic clades, while the incompatible strains belonged to separate phylogenetic clades (Fig. 1C). These results indicate that the phylogenetic distance between *A. oryzae* strains is important in determining compatibility.

### Compatibility analysis via natural cell fusion in co-culture

Owing to the forced protoplast fusion in the auxotrophic complementation assay, it was still unclear whether strains can distinguish between compatible and incompatible pairings. To investigate the natural cell fusion ability of compatible/incompatible pairings, we performed compatibility analysis using a co-culture assay, where strains with *pyrF* and *pyrG* deletions were allowed to naturally undergo cell fusion (Fig. 2A). Two different auxotrophic strains Δ*pyrF* and Δ*pyrG* were co-cultured for 5 days in a minimal medium supplemented with uridine/uracil, and the collected conidia were spread on minimal medium without supplementation. Only heterokaryon conidia, formed via cell fusion with auxotrophic complementation, could grow in the minimal medium. The cell fusion efficiency was then quantified by counting the colony number, and thus, the compatible/incompatible relationships could be evaluated based on growth.

**FIG 2.**
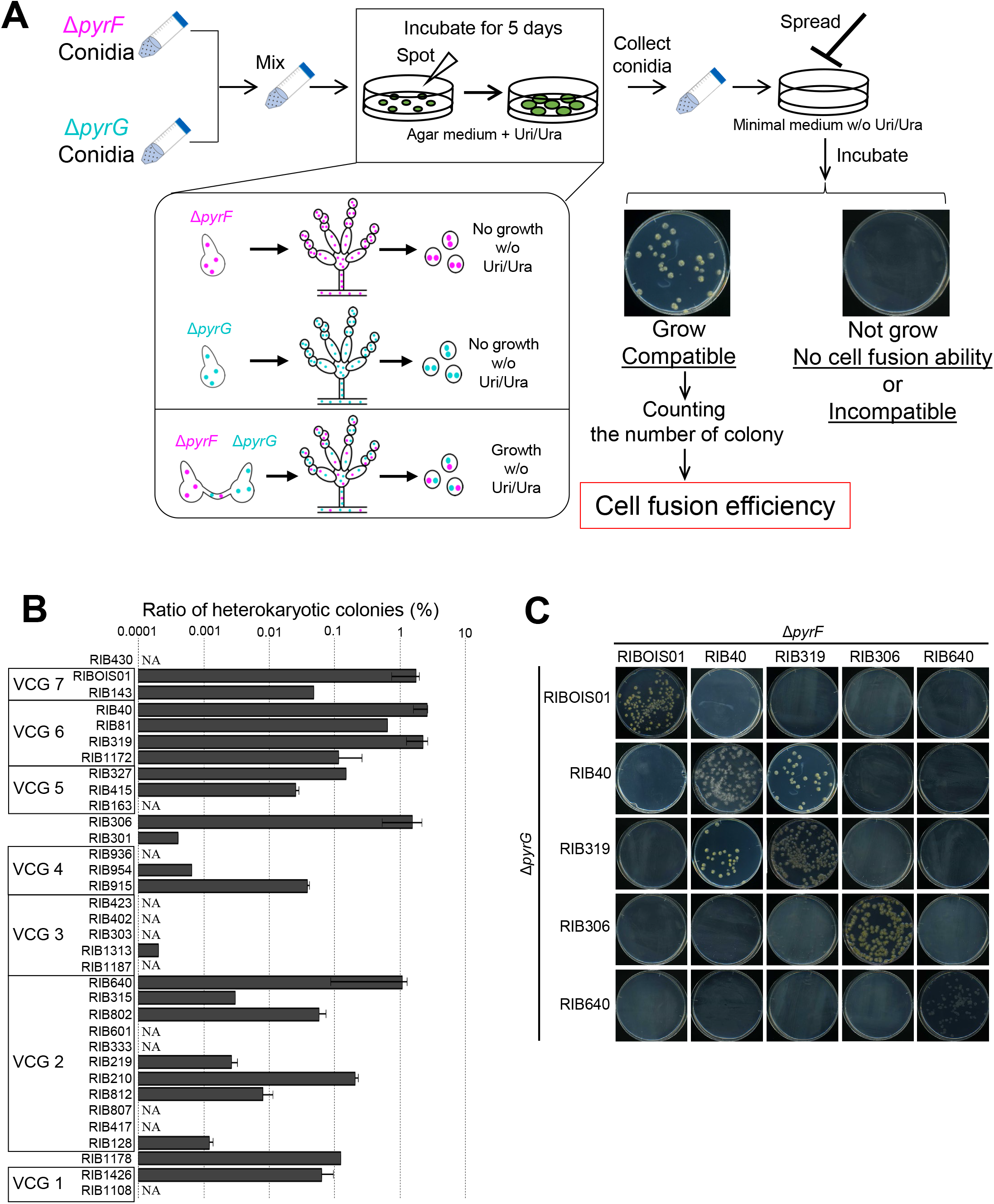
Heterokaryon incompatibility analysis via co-culture assay. (A) Protocol of co-culture assay. Equal numbers of conidia from two uridine/uracil auxotrophic strains (Δ*pyrF* and Δ*pyrG*) were mixed, and 5 × 10^4^ conidia/5 μL were spotted onto the agar medium containing uridine and uracil. After incubation at 30°C for 5 days, the newly formed conidia were collected, and spread on the minimal agar medium without uridine and uracil. Only heterokaryotic conidia auxotrophically complemented grew in the CD minimal medium, and the colony size was restricted by the addition of 0.25% Triton X-100. By counting the number of colonies, cell fusion efficiency was quantified. (B) Quantification of the self-fusion efficiency in 34 *A. oryzae* strains. The error bars indicate standard deviations based on Student’s *t* tests. Three independent experiments were performed. NA: Not analyzed. (C) Incompatibility analysis via co-culture assay involving five *A. oryzae* strains with higher self-fusion efficiency.

As 10 out of the 34 selected strains did not form enough conidia, self-fusion efficiency was quantified using 24 strains (Fig. 2B). Five strains showed higher self-fusion efficiencies with a heterokaryon conidia ratio above 1%. The use of strains with higher fusion efficiencies can facilitate the investigation of heterokaryon formation via natural cell fusion, and therefore, the five strains with higher self-fusion efficiency were analyzed for VCG classification. We found that only the strains RIB40 and RIB319 were compatible (Fig. 2C), fully consistent with the observations when forced protoplast fusion was performed (Fig. 1AB), which further supported the reliability of our VCG classification via auxotrophic complementation assay (Fig. 1C).

### Visualization of physiological responses in incompatible strain pairs

To investigate how strains distinguish between compatible/incompatible ones, the visualization of physiological responses at cellular level was necessary. Additionally, we suspected that the use of strains with higher self-fusion efficiency would facilitate cellular observation. Thus, the strains RIB40 and RIB640, that showed higher self-fusion efficiencies, were selected and their cytoplasm were labelled with fluorescent proteins using the CRISPR/Cas9 system (Fig. S3A). After mixing the conidia derived from the strain pair, the mixture was spotted onto the optimized minimal medium to enhance the cell fusion efficiency (24), which was later analyzed for time-course fluorescence observations (Fig. 3A).

**FIG 3.**
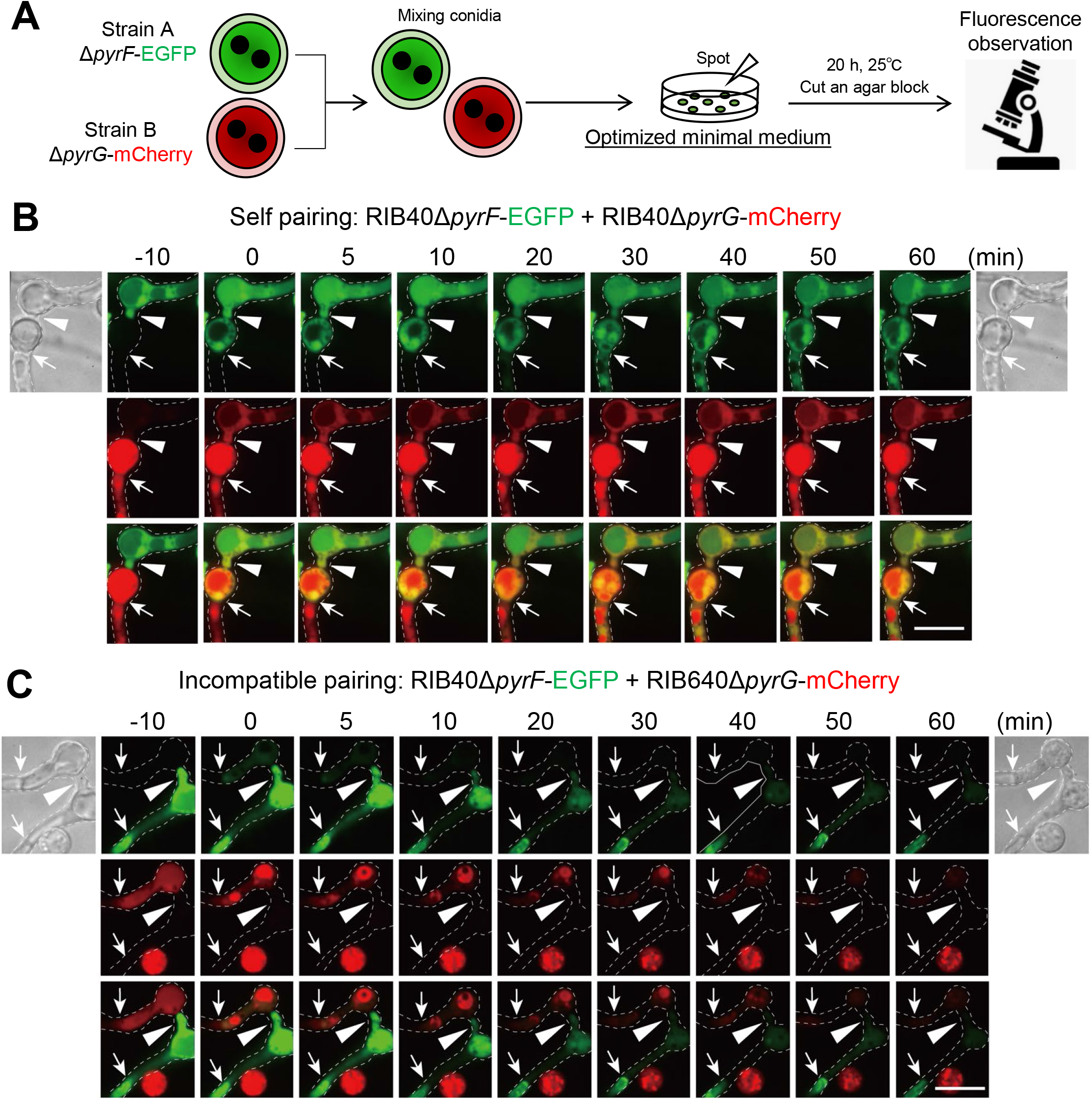
Visualization of physiological responses to incompatible cell fusion in the cytoplasm. (A) The time-course fluorescence observation method. (B) Gradual cytoplasmic mixing of EGFP and mCherry in heterokaryotic cells following the self-pairing RIB40Δ*pyrF*-EGFP and RIB40Δ*pyrG*-mCherry strains. Arrowheads and white arrows indicate fusion points and septa, respectively. Scale bar represents 10 µm. (C) Cytoplasmic fluorescence loss in the fused area of incompatible pairing between RIB40Δ*pyrF*-EGFP and RIB640Δ*pyrG*-mCherry strains. Note that cell fusion started at the 0-min time point as indicated by the entry of EGFP fluorescence into the cell of the mCherry-labelled strain. Arrowheads and white arrows indicate fusion points and septa, respectively. Scale bar represents 10 µm.

For self-pairings, two fluorescence proteins were first mixed only in the fused cells, while the septum of the adjacent cell was closed (Figs. 3B and S3B). Within 20–30 min after cell fusion, the cytoplasmic fluorescence was transferred into the next cell by opening the septum (Figs. 3B and S3B). During incompatible pairing, cell fusion also occurred initially as detected by the slight entrance of EGFP into the mCherry-labeled cells (Fig. 3C). This was followed by the gradual loss of fluorescence in the fused cell, while septum closure maintained the fluorescence in the adjacent cell (Fig. 3C).

To characterize the fluorescence loss observed during the fusion of incompatible cells, the cell death indicator, propidium iodide (PI), a red fluorescent dye that only stains the nuclei of dead cells, was added to the incompatible pairing co-culture. We did not observe PI staining during the entire self/compatible cell fusion process (Fig. 4A). However, during incompatible pairing, PI-stained nuclei appeared in the fusion area within 10–20 min after cell fusion, while the cells across the septum did not show any staining (Fig. 4B). Simultaneously, EGFP fluorescence in the fusion area showed a gradually decreasing trend, indicating that cell death occurred rapidly during incompatible cell fusion. As elevated ROS levels are associated with incompatibility-triggered cell death (21), we also examined ROS accumulation using the oxidant-sensitive probe, 2’,7’-dichlorodihydrofluorescein diacetate (H_2_DCFDA), which fluoresces via ROS-mediated oxidization. We observed significant ROS accumulation in the fused area corresponding to incompatible pairing, whereas self-pairing showed no fluorescence (Figs. 4C and S3C).

**FIG 4.**
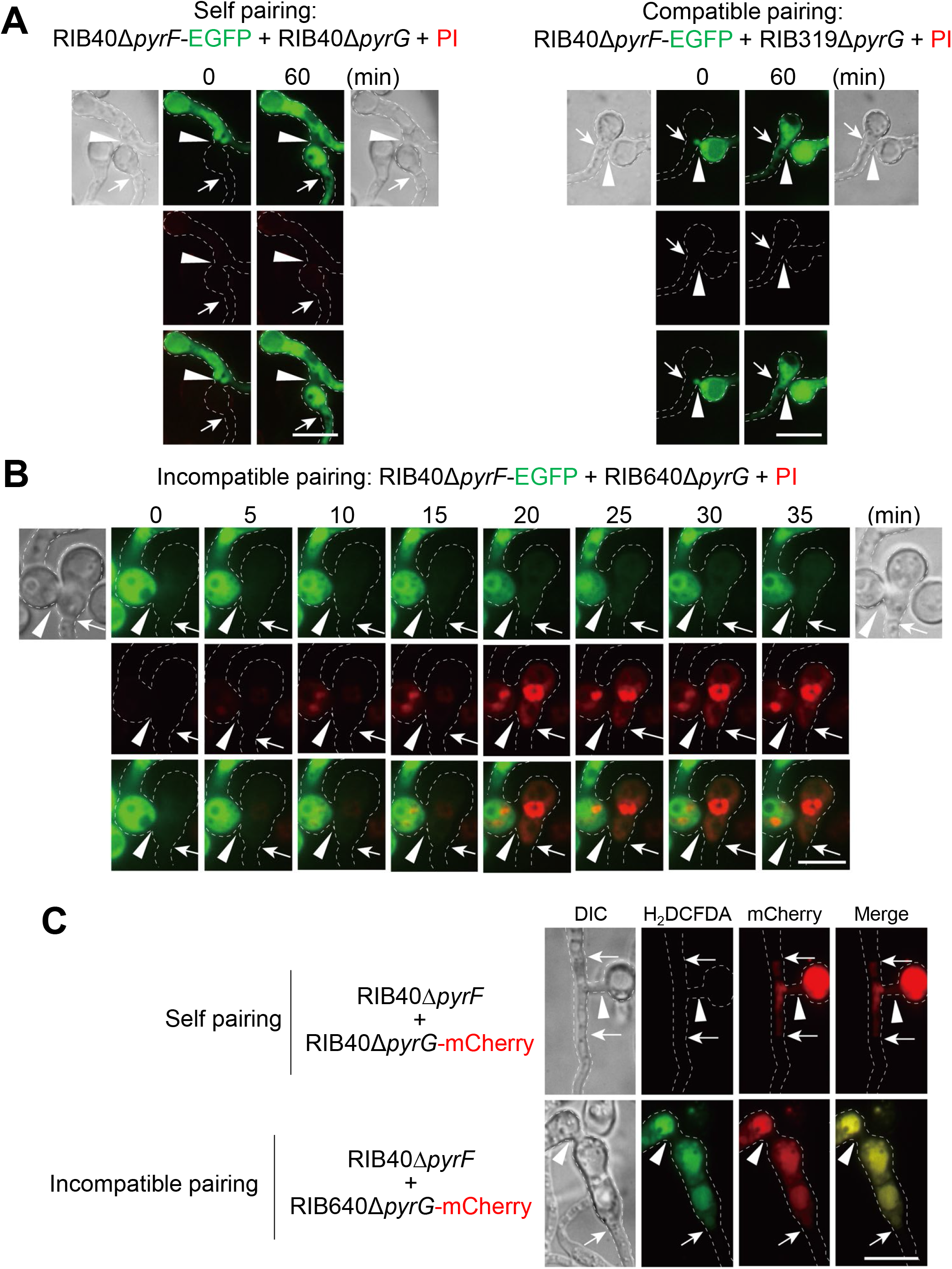
Incompatibility-triggered cell death following cell fusion. (A) PI staining in self/compatible pairing between RIB40 and RIB319 strains during the fusion process. Arrowheads and white arrows indicate fusion points and septa, respectively. Scale bars represent 10 µm. (B) PI staining in incompatible pairing between RIB40 and RIB640 strains. Note that cell fusion started at the 0-min time point as indicated by the entry of EGFP fluorescence into the cells of the non-labelled strain. Arrowheads and white arrows indicate fusion points and septa, respectively. Scale bar represents 10 µm. (C) ROS accumulation in the fused area of self/incompatible pairings between RIB40 and RIB640 strains. Arrowheads and white arrows indicate fusion points and septa, respectively. Scale bar represents 10 µm.

To further characterize the subcellular dynamics associated with incompatible responses, we visualized organelles such as the nuclei, endoplasmic reticulum (ER), and mitochondria. To test the incompatible effect on nuclei, the gene necessary for expressing the histone H1-mCherry fusion protein was introduced into the strains RIB40Δ*pyrG* and RIB640Δ*pyrG*, yielding RIB40Δ*pyrG-*H1-mCherry and RIB640Δ*pyrG-*H1-mCherry (Fig. S3D), respectively, which were later paired with the strain RIB40Δ*pyrF-*EGFP. After cell fusion of self-pairing, indicated by the entry of EGFP fluorescence into the mCherry-labeled strain, visible red fluorescence and the round shape of the nuclei were maintained (Fig. S4AB). However, during incompatible pairing, histone H1-mCherry fluorescence was gradually lost after cell fusion, as indicated by the entry of EGFP fluorescence into the paired strain (Fig. S4C). The mCherry-labeled nuclei in the adjacent cell were protected, probably due to septum closure, as the red fluorescence across the septum was retained. For ER visualization, we used the chaperone BiP, and the *AobipA-egfp* fusion gene was introduced into the strains RIB40Δ*pyrF* and RIB640Δ*pyrF* (Fig. S3D), which were later paired with the strain RIB40Δ*pyrG-m*Cherry. When mCherry fluorescence entered the AoBipA-EGFP-expressing strain during self-pairing, AoBipA-EGFP fluorescence did not significantly change, and the ER-tubular network remained intact (Fig. S5AB). In contrast, after incompatible cell fusion, indicated by the entry of mCherry fluorescence into the cell of the AoBipA-EGFP-expressing strain, AoBipA-EGFP fluorescence gradually decreased (Fig. S5C). These findings suggest that incompatible physiological responses have disruptive effects on the nuclei and ER.

The effects of incompatible responses on mitochondria were also investigated. The mitochondrial protein marker, mitochondrial citrate synthase (AoCit1), fused with EGFP was used to yield the RIB40Δ*pyrF-*AoCit1-EGFP and RIB640Δ*pyrF-*AoCit1-EGFP strains (Fig. S3D). Compared with the mitochondrial morphology in self-pairing (Figs. 5A and S6A), the tubular structure of the mitochondria associated with incompatible fusion became punctuate immediately after the incompatible cells fused, and the fluorescence of the fragmented mitochondria gradually decreased (Fig. 5B). Furthermore, fragmented mitochondria also appeared in the swapped incompatible strain pairs during the cell fusion process (Fig. S6B). Thus, heterokaryon incompatibility directly resulted in mitochondrial fragmentation.

**FIG 5.**
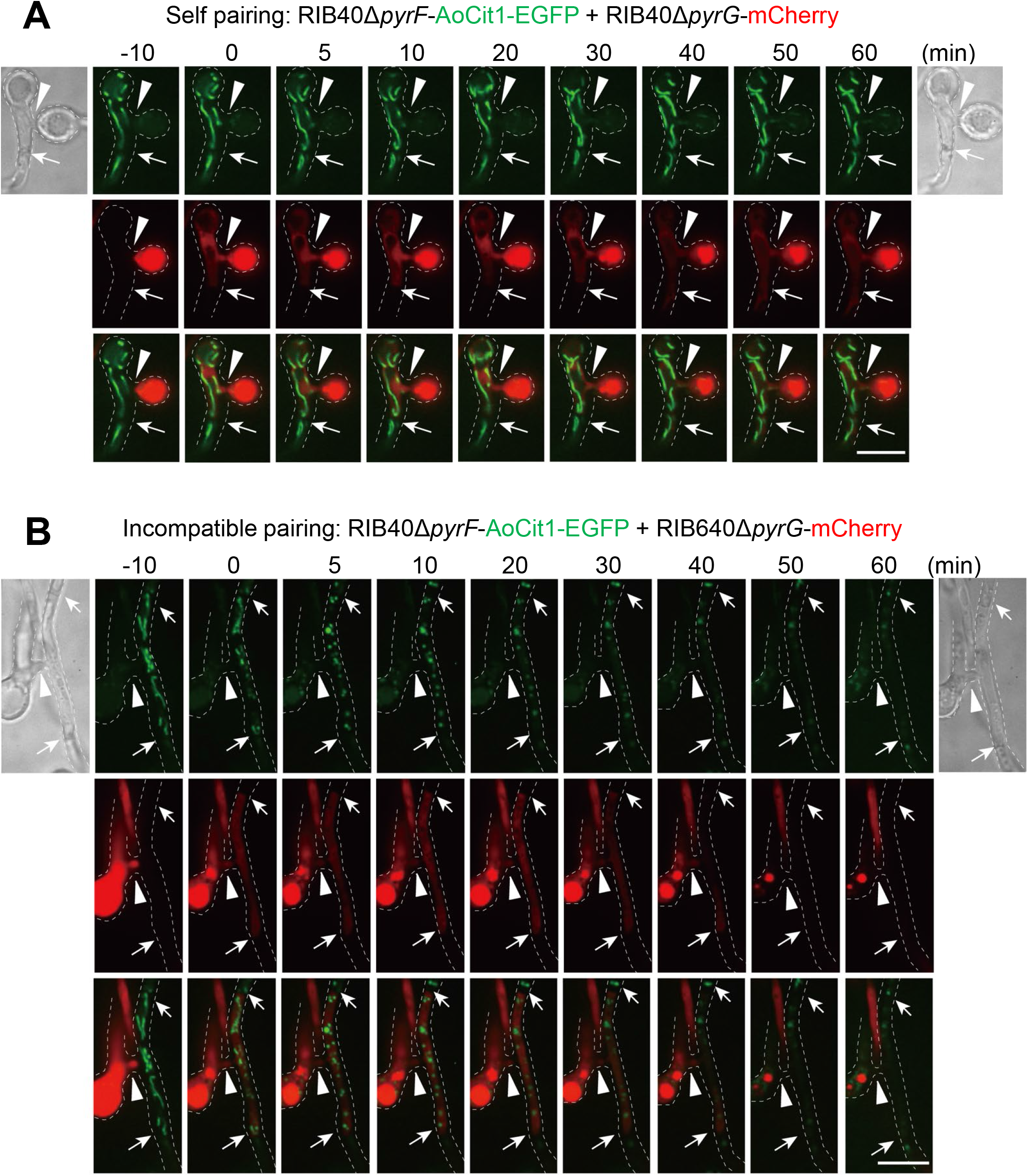
Visualization of incompatibility-induced physiological responses in mitochondria. (A) Gradual entry of mCherry fluorescence into the cell of the AoCit1-EGFP-labelled strain during RIB40Δ*pyrF*-EGFP and RIB40Δ*pyrG*-mCherry self-pairing. Arrowheads and white arrows indicate fusion points and septa, respectively. Scale bar represents 10 µm. (B) AoCit1-EGFP-labeled mitochondrial fragmentation and consequent fluorescence loss after the cell fusion of the incompatible strain pair, RIB40Δ*pyrF*-EGFP and RIB640Δ*pyrG*-mCherry. Arrowheads and white arrows indicate fusion points and septa, respectively. Scale bar represents 10 µm. Note that cell fusion started at the 0-min time point as indicated by the entry of mCherry fluorescence into the cells of the AoCit1-EGFP-labelled strain.

### Effect of the deletion of mitochondrial fission-related genes on incompatibility-triggered cell death

The characteristics of mitochondrial fragmentation following incompatible cell fusion suggested that the mitochondrial fission machinery may be an important mechanism that regulates the fate of heterokaryotic cells. Reportedly, mitochondria are dynamic organelles that undergo constant fusion and fission during normal cell division, and the equilibrium between fission and fusion is controlled by the activity of conserved molecular machineries driven by dynamin-like GTPases (25). The dynamin-related protein Dnm1 and mitochondrial fission 1 Fis1 are the main regulators of mitochondrial fission in eukaryotic organisms (26).

To investigate the correlation between changes in mitochondrial morphology and incompatibility-triggered cell death, we analyzed the *A. oryzae* genome for the orthologs of mitochondrial fission-related genes. We selected *Aofis1* (AO090026000294) and *Aodnm1* (AO090010000776) for deletion (Fig. S7A). To examine whether mitochondrial fission deficiency blocks incompatibility, leading to the formation of the heterokaryon on a minimal medium, we performed a protoplast fusion assay on the strain pairing with the gene deletions. Even though all the incompatible pairings with the deletions did not produce any clearly visible mycelia (Fig. S7B), heterokaryon hyphae were found using a microscope (Fig. 6A). Similar to the control, the recovered heterokaryons did not show any PI staining, suggesting viability (Fig. S7C). Additionally, the quantification of the number of grown heterokaryons in incompatible pairing with gene deletions clearly indicated the recovery of heterokaryons in incompatible pairings with *Aodnm1* and *Aofis1* deletions (Fig. 6B).

**FIG 6.**
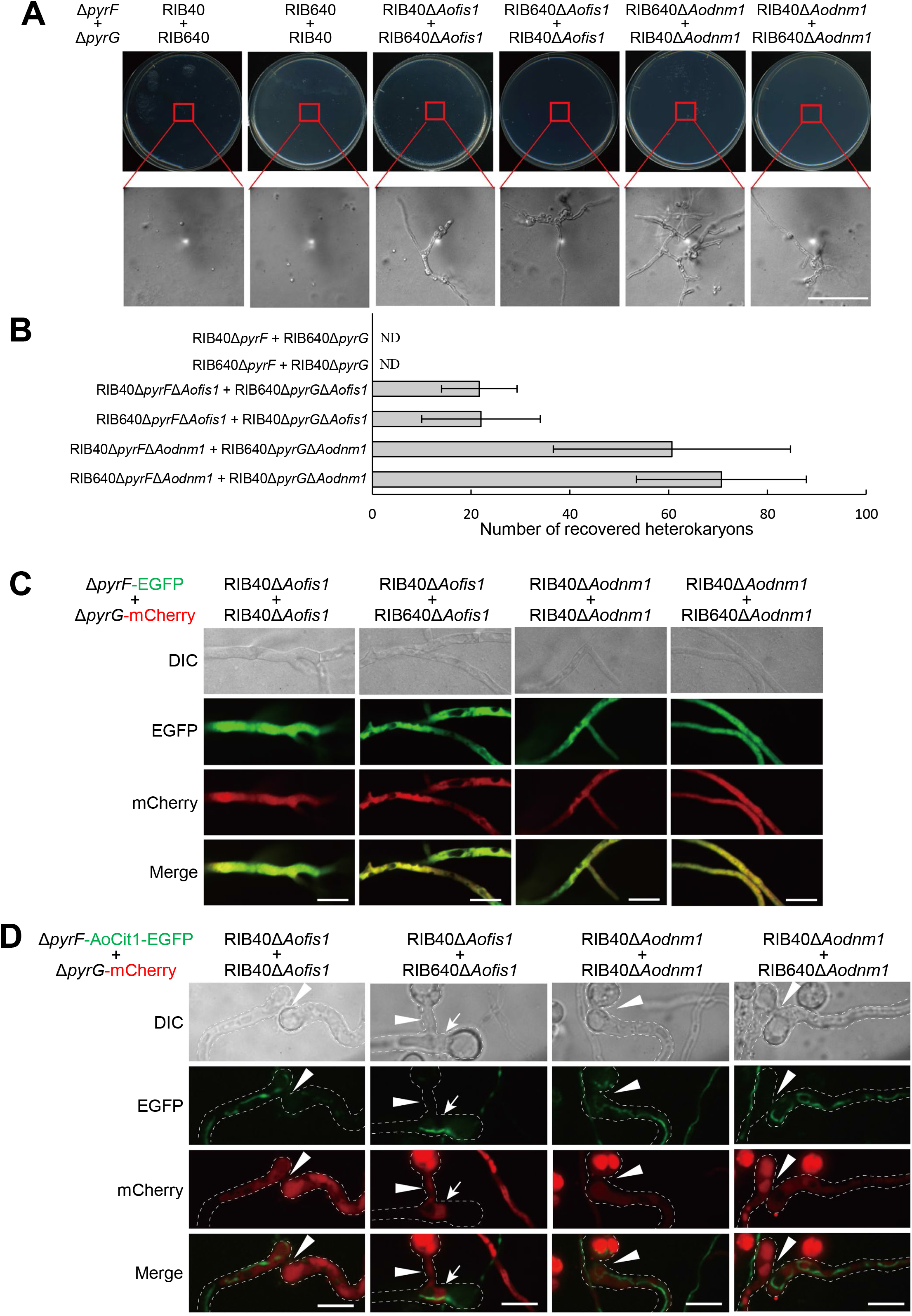
Alleviation of incompatibility-triggered cell death by deletion of mitochondrial fission-related genes. (A) Microscopic observation of recovered heterokaryon following the deletion of *Aofis1* and *Aodnm1* in incompatible pairing. Protoplasts in non-self-pairing with different uridine/uracil auxotrophies (Δ*pyrF* and Δ*pyrG*) were forcedly fused using polyethylene glycol (PEG). The fused protoplasts were plated on the CD minimal agar medium without uridine/uracil at 30°C for 5 days. Scale bar represents 100 µm. (B) Quantification of the number of heterokaryons in the different pairings. The error bars indicate standard deviations. Three independent experiments were performed. ND: Not detected. (C) Recovered heterokaryons with mixing cytoplasmic fluorescence, indicating that they were derived from the forced protoplast fusion of incompatible pairing with *Aofis1* and *Aodnm1* deletions. Scale bars represent 10 µm. (D) Retention of the tubular structure of mitochondria after the deletion of mitochondria fission-related genes in incompatible cell fusion during the co-culture assay. Arrowheads and white arrows indicate fusion points and septa, respectively. Scale bars represent 10 µm.

To validate the nature of the heterokaryons grown during the incompatible pairing of the strains with *Aodnm1* and *Aofis1* deletions, we labelled each paired strain with different fluorescence (Fig. S8A), which were later applied to the protoplast fusion assay. The recovered heterokaryons in incompatible pairing with *Aodnm1* and *Aofis1* deletions showed mixed cytoplasmic fluorescence, which was comparable with the result observed for compatible pairings (Fig. 6C). This observation indicated that the recovered heterokaryons were derived from incompatible cell fusion. Further, to visualize whether mitochondrial fragmentation still occurred during incompatible pairing involving the strains with *Aodnm1* and *Aofis1* deletions, we constructed the strains RIB40Δ*pyrF*Δ*Aofis1*-AoCit1-EGFP and RIB40Δ*pyrF*Δ*Aodnm1*-AoCit1-EGFP and then co-cultured them with the corresponding strains to visualize mitochondrial morphology (Fig. S8B). The fused cells in incompatible pairing involving the strains with *Aodnm1* and *Aofis1* deletions showed tubular-structured mitochondria similar to those in compatible pairings (Fig. 6D). These observations indicate that incompatibility-triggered cell death is successfully alleviated following the deletion of mitochondrial fission-related genes *Aofis1* and *Aodnm1*, demonstrating that mitochondrial fragmentation has physiological relevance in incompatibility responses.

## DISCUSSION

Determining whether members of the same VCG originate from the same ancestor is of interest. In a previous study, phylogenetic analysis involving the VCGs of *Verticillium dahlia* suggested that isolates belonging to the same group have similar sets of nucleotide polymorphisms markers and DNA sequences in the specific genomic regions (27). Using a species-specific DNA probe, it was observed that multiple *A. flavus* strains belonging to the same VCG display identical DNA fingerprints (28). This observation is also supported by a comparison of VCGs, which involved the analyses of random amplified polymorphic DNA (29) and single nucleotide polymorphisms of the specific genes (30). However, evidence at the overall strain level has not yet been elaborated. In this study, 34 *A. oryzae* strains, which comprehensively represent all the phylogenetic clades of this fungus, were subjected to an auxotrophic complementation assay; at least seven VCGs were identified, and it was also observed that strains of the same group belonged to the same or phylogenetically close clades, while incompatible strains were phylogenetically distant (Fig. 1C). Thus, we concluded that strains belonging to the same VCG show genetic composition similarities.

Cell fusion between different individuals and the establishment of viable heterokaryons are believed to be largely prevented by heterokaryon incompatibility. To track this cell fusion process at cellular level, the discrimination of individual cells using different fluorescence proteins or staining dyes is a useful strategy; in *N. crassa*, one cell expresses cytoplasmic GFP, while the other paired cell is stained with FM4-64 dye (31). Owing to the low cell fusion efficiency of *A. oryzae*, dynamic observation of cell fusion events is difficult. However, in this study, we selected strains with higher self-fusion efficiency (Fig. 2B) and used them in a co-culture assay. The compatibility/incompatibility results in the co-culture assay (Fig. 2C) were consistent with those in the forced protoplast fusion (Fig. 1AB). These observations enhanced the authenticity of our VCG classification (Fig. 1C). Additionally, starvation conditions significantly induce the cell fusion of the fungus *V. dahlia* (32), and media components have been optimized for efficient *A. oryzae* fusion (24). These previous studies motivated us to eliminate nitrogen sources and increase glucose concentration to 20% in the optimized minimal medium, which further enhanced the fusion efficiency of *A. oryzae* self-pairing, as well as that of non-self-pairing. These optimizations allowed us to perform time-course fluorescence observations to capture cell fusion event processes in two fluorescence-labeled *A. oryzae* self-pairings (Figs. 3B and S3B).

More importantly, the process of incompatibility-triggered cell death was successfully visualized as directly evidenced by PI staining (Fig. 4B). Our results also indicated that incompatibility-triggered cell death involved various physiological responses including ROS accumulation (Fig. 4C) and septal plugging (Fig. 3C), which are typically observed in heterokaryon incompatibility (17–21, 33). Further, time-course observations of physiological responses based on PI staining indicated that cell death started within 10–20 min after incompatible cell fusion (Fig. 4B). Subsequently, the gradual disappearance of fluorescence markers for cytosol, nuclei, ER, and mitochondria was also observed (Figs. 3C, 5B, S4C, S5C, and S6B). We assumed that indiscriminate protein degradation or fluorescent protein diffusion possibly occurred as secondary events owing to incompatibility-triggered cell death.

Notably, the most significant physiological response to incompatible cell fusion was immediate mitochondrial fragmentation (Fig. S6B). Moreover, this incompatibility-triggered cell death was alleviated when strains with *Aodnm1* and *Aofis1* deletions were paired (Fig. 6). Thus, this study is the first to demonstrate that mitochondrial fission-related proteins are involved in heterokaryon incompatibility-triggered cell death (Fig. 7). Generally, mitochondria are considered to play a central role in programmed cell death in fungi. In *Saccharomyces cerevisiae*, it has been observed that the loss of the mitochondrial fission machinery leads to resistance to acetic acid-induced cell death (34). In *Aspergillus fumigatus*, mitochondrial fragmentation was found to be associated with azole-induced cell death (35), and the loss of the mitochondrial fission machinery resulted in azole resistance (36). In the filamentous ascomycete *P. anserina*, a functional link of mitochondrial fission was found to be associated with apoptosis and life-span control (37). Therefore, the importance of mitochondrial fission in inducing cell death is a phenomenon common to various cell death-related processes in yeasts and filamentous fungi. However, the deletion of mitochondrial fission-related genes did not completely suppress heterokaryon incompatibility in *A. oryzae* (Fig. 6A), indicating the possibility that other factors/pathway are involved in cell death (Fig. 7). It has been suggested that in *N. crassa*, two parallel pathways with VIB-1 and protein kinase IME-2 are responsible for the regulation of heterokaryon incompatibility and cell death, and mitochondrial involvement in cell death via IME-2 was proposed (38). Further investigation of the multiple machineries including mitochondrial fission would uncover the general mechanisms governing heterokaryon incompatibility-triggered cell death. It is also expected that the findings of such investigations will help overcome the incompatibility barrier that limits the industrial crossbreeding of filamentous fungi.

**FIG 7.**
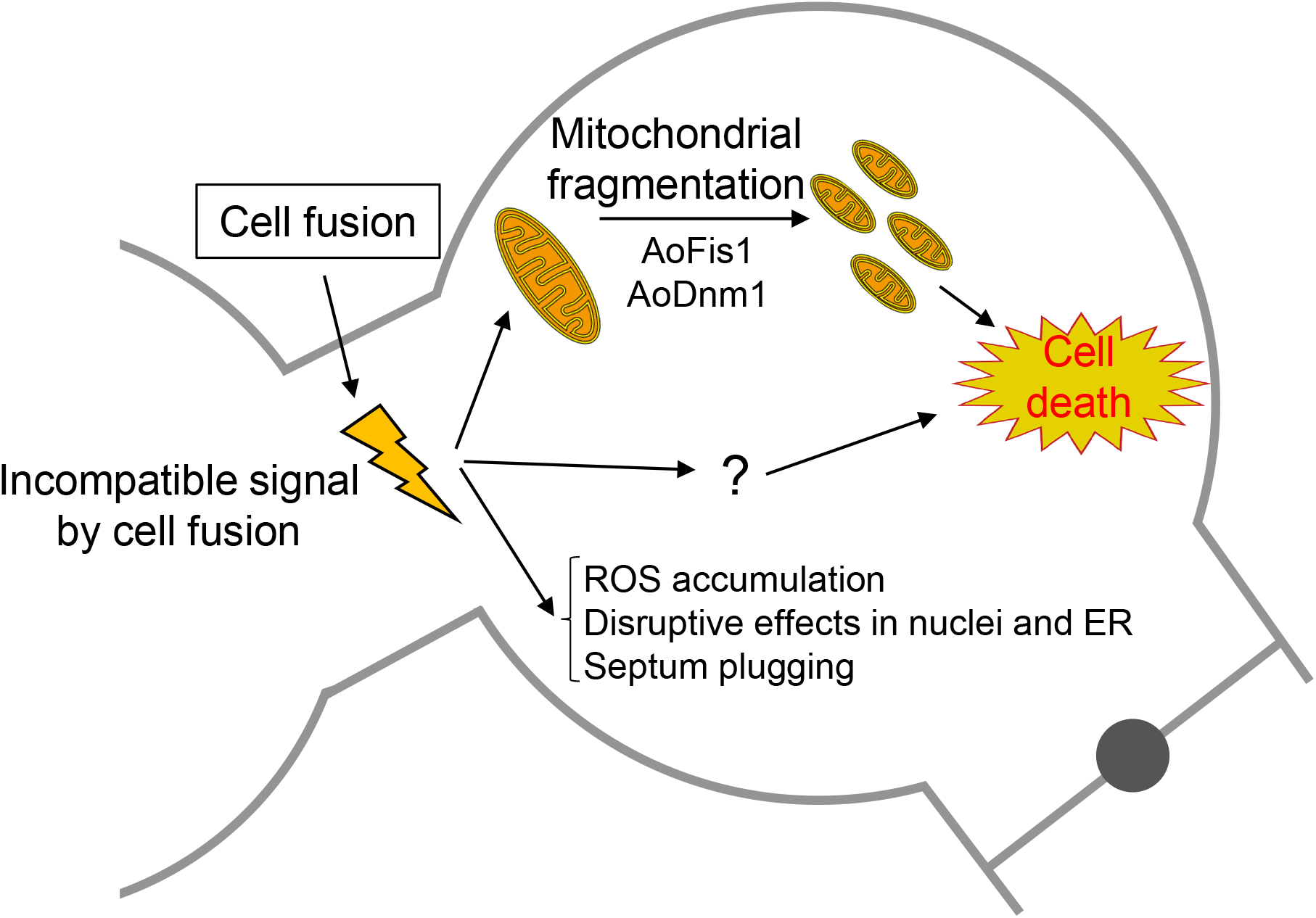
A proposed model of the mechanism underlying the regulation of incompatibility-triggered cell death by mitochondrial fission-related proteins AoFis1 and AoDnm1. When cells received an incompatibility signal induced by incompatible cell fusion, various physiological responses such as mitochondrial fragmentation were observed in different organelles. Two mitochondrial fission-related proteins AoFis1 and AoDnm1 were found to promote the cell death process.

## MATERIALS AND METHODS

### *A. oryzae* strains, growth conditions, and transformation

The *A. oryzae* strains used in this study, which are listed in Table S1, were grown in Czapek-Dox (CD) minimal medium (0.3% NaNO_3_, 0.2% KCl, 0.1% KH_2_PO_4_, 0.05% MgSO_4_·7H_2_O, 0.002% FeSO_4_·7H_2_O, and 2% glucose; pH 5.5) and potato dextrose (PD) medium (Nissui, Tokyo, Japan). DPY medium (2% dextrin, 1% polypeptone, 0.5% yeast extract, 0.5% KH_2_PO_4_, and 0.05% MgSO_4_·7H_2_O; pH 5.5) was used for pre-culturing during transformation. For selection using the *ptrA* marker, the DPY medium without yeast extract (DP) was used for pre-culture. To select the Δ*pyrF* and Δ*pyrG* strains, 5-fluoroorotic acid (5-FOA) was added to the medium, and to grow these selected strains, 0.2% uridine and 0.1% uracil were added. For fluorescence microscopic analysis, the strains were grown in an optimized minimal medium (0.2% KCl, 0.1% KH_2_PO_4_, 0.05% MgSO_4_·7H_2_O, 0.002% FeSO_4_·7H2O, 20% glucose, and 0.005% uridine; pH 7.0).

In this study, strains were constructed by transformation using the CRISPR/Cas9 system, in which the genome editing plasmid and the donor plasmid were used for co-transformation. The transformation of *A. oryzae* was performed as previously described (39) with some modifications. The genome-editing plasmid and the donor plasmid (ratio 3:5) were used for co-transformation with the protoplast suspension and incubated on ice for 15 min. Polyethylene glycol solution (60% PEG4000, 50 mM CaCl_2_, and 10 mM Tris-HCl; pH 7.5) was added twice (first 500 μL and then 800 μL), and after incubation at 25°C for 20 min, the PEG-treated protoplast suspension was mixed with the top agar selective medium, and immediately spread on CD agar medium. For deletion of *pyrF*/*pyrG* and introduction of fluorescent protein expression cassettes, CD medium containing 0.1 ug/mL pyrithiamine was used as the top agar medium. For *Aofis1/Aodnm1* deletions, *pyrF* and *pyrG* were used as selection markers, and CD medium was used as the top agar medium.

### Plasmid construction

The primers used for plasmid construction are listed in Table S2. DNA fragments were amplified from the genomic DNA of RIB40 using PrimeSTAR^®^ HS DNA Polymerase (TaKaRa, Otsu, Japan) and ligated with vectors using the In-Fusion^®^ HD Cloning Kit (TaKaRa).

To delete *pyrF/pyrG* genes in 34 *A. oryzae* strains, pRGE-pyrF/ppAsATC9a2gpG and pUC-pyrF-donor/pUC-pyrG-donor were used as the genome editing and donor plasmids, respectively. To construct the pRGE-pyrF plasmid, the *U6* promoter (P*U6*) with the target sequence to *pyrF* was amplified from the pRGE-gwAup plasmid (40) and ligated with the *Sma*I-digested pRGE-gRT6 plasmid. To construct the ppAsATC9a2gpG plasmid, P*U6* with target sequence to *pyrG* and the *U6* terminator (T*U6*) with the scaffold sequence of sgRNA were amplified from pRGE-gwAup and then ligated with the *Sma*I-digested ppAsATC9a2 (40). To construct the pUC-pyrF-donor and pUC-pyrG-donor plasmids, the upstream and downstream fragments of *pyrF*/*pyrG*, respectively, were ligated with the *Bam*HI-digested pUC19 plasmid.

To investigate the expression of cytoplasmic fluorescence proteins in the Δ*pyrF/pyrG* strain background, the genome editing plasmid pRGE-gwAup and the donor plasmids were co-introduced. To construct the donor plasmids pwAup-EGFP-tef1PT and pwAup-mCherry-tef1PT, the *tef1* promotor/terminator of *A. oryzae* was amplified and ligated with the *Sma*I-digested pwAupDN plasmid (40), yielding pwAup-tef1PT. Additionally, *egfp* and *mcherry* fragments were amplified and ligated with the *Sma*I-digested pwAup-tef1PT plasmid.

To construct strains expressing organelle marker fluorescence proteins, donor plasmids were constructed. To construct the donor plasmid pwAup-PH1-H1-mCherry, the DNA fragment containing the mCherry-encoding gene and the *amyB* terminator (T*amyB*) amplified from pisCIIA-mCherry (41) were inserted into the *Sma*I site of pisC (41), yielding pisC-C-mCherry. The DNA fragment containing the promoter and ORF of the histone H1 gene (AO090701001235) was ligated with the *Sma*I-digested pisC-C-mCherry plasmid, yielding pisC-h1mC, while the P*h1*::*h1-mCherry*::T*amyB* fragment was amplified from the pisC-H1-mC plasmid and ligated with the *Sma*I-digested pwAupDN plasmid. To construct the donor plasmid pwAup-BipA-EGFP-amyBPT, the P*amyB*::*AobipA-egfp*::T*amyB* fragment was amplified from the pUNABG plasmid (42), and ligated with the *Sma*I-digested pwAupDN plasmid. To construct the donor plasmid pwAup-Cit1-EGFP-amyBPT, the fragment of *Aocit1* was amplified from the pgACEN plasmid (43) and ligated with the *Sma*I-digested pwAupEGFP plasmid (40).

For *Aofis1*/*Aodnm1* deletions, genome editing plasmids and donor plasmids were used for co-transformation. The pRpG-gRT6 plasmid, which was used for the construction of the genome editing plasmid, was constructed as follows: the *pyrG* gene was amplified and ligated with the *Sma*I-/*Bst*1107I-digested ppAsAMA plasmid (40), yielding ppGAMA. The *cas9*-expression cassette with P*tef1* and T*amyB*, amplified from ppAsATC9 (40), was inserted into the *Hin*dIII site of ppGAMA, yielding the ppGsATC9 plasmid. Then, the T*U6* with the scaffold sequence of sgRNA was amplified from pRGE-gRT6, yielding pRpG-gRT6. The P*U6* fragment with the target sequence was amplified from the pRGE-gwAup plasmid and ligated with the *Sma*I-digested pRpG-gRT6 plasmid. The ppFsATC9-gRT6 plasmid, which was used for the construction of the genome editing plasmid, was constructed as follows: the amplified *pyrF* fragment was ligated with the *Sma*I-*Bst*1107I digested ppGsATC9 plasmid, yielding ppFsATC9. Then, the T*U6* with the scaffold sequence of sgRNA was amplified from pRGE-gRT6, yielding ppFsATC9-gRT6. The P*U6* fragment with the target sequence was amplified from the pRGE-gwAup plasmid and ligated with the *Sma*I-digested ppFsATC9-gRT6 plasmid.

For *Aofis1* deletion, the genome editing plasmids ppFsATC9-gFis1 and pRpG-gFis1 were constructed as follows: The P*U6* with the sgRNA sequence targeting the *Aofis1* gene was amplified from the pRGE-gwAup plasmid and ligated with the *Sma*I-digested ppFsATC9-gRT9 and pRpG-gRT6 plasmids. To construct the donor plasmid, pFis1-donor, the upstream or downstream fragments of the *Aofis1* ORF were amplified and ligated with the *Bam*HI-digested pUC19 plasmid.

For *Aodnm1* deletion, the genome editing plasmids ppFsATC9-gDnm1 and pRpG-gDnm1 were constructed as follows: P*U6* with the sgRNA sequence targeting the *Aodnm1* gene was amplified from the pRGE-gwAup plasmid and ligated with the *Sma*I-digested ppFsATC9-gRT9 and pRpG-gRT6 plasmids. To construct the donor plasmid, pDnm1-donor, the upstream or downstream fragments of the *Aodnm1* ORF were amplified and ligated with the *Bam*HI-digested pUC19 plasmid.

### Protoplast fusion assay

Based on the *A. oryzae* transformation method without DNA introduction, a protoplast fusion assay was performed with some modifications to the protocol. To classify the VCGs, the following protoplast fusion assay was performed. Equal volumes (200 μL) of the protoplast suspension (1 × 10^7^/mL) obtained from two different auxotrophic strains were mixed, and then, a PEG solution (60% PEG4000, 50 mM CaCl_2_, and 10 mM Tris-HCl; pH 7.5) was added twice (first 500 μL and then 800 μL). After incubation at room temperature for 20 min, CD medium containing 0.2% uridine and 0.1% uracil was added into the protoplast suspension, which was then immediately spread on CD agar medium containing 1.2 M sorbitol without uridine or uracil. After incubation at 30°C for 3 days, the agar plates were inspected for the growth of hyphae (heterokaryon).

### Co-culture assay

To investigate strains with natural cell fusion ability, the following co-culture protocol was applied: conidia of two different auxotrophic strains (Δ*pyrF* and Δ*pyrG*) were collected from PD agar medium containing uridine/uracil. Equal volumes of conidial suspension (1 × 10^7^ conidia/mL) from both strains were mixed, and 5 μL aliquots were spotted onto the CD agar media containing uridine/uracil. After incubation at 30°C for 5 days, the newly formed conidia were collected, and 1 × 10^4^–1 × 10^6^ conidia were spread onto a selective agar medium without uridine/uracil. To accurately count the number of colonies, 0.25% Triton X-100 was added to the selective agar medium to reduce colony size. After incubation, the colonies derived from auxotrophically complemented heterokaryons were counted. Additionally, to observe fluorescence, equal numbers of conidia from corresponding fluorescent strains were mixed, and 5 μL aliquots of the conidial suspension (1 × 10^7^ conidia/mL) were spotted onto the optimized minimal media for incubation at 25°C for 20 h.

### Fluorescence microscopic analysis

Equal aliquots of conidial suspension (1.0 × 10^7^ conidia/mL) from self/incompatible strain pairs were mixed and spotted on the optimized minimal medium. Different strain pairs were either discriminated by EGFP vs. mCherry, EGFP vs. no fluorescence, or EGFP vs. PI (3.33 μg/mL) staining. The agar plates were then incubated for 20 h at 25°C, and agar squares (1 cm^2^) were excised and turned upside down on a glass bottom dish for fluorescence observation. The cultures were observed by confocal laser scanning microscopy using an IX71 inverted microscope (Olympus, Tokyo, Japan), with 488-nm (Furukawa Electric, Tokyo, Japan) and 561-nm (Melles Griot, Rochester, NY, USA) semiconductor lasers, EGFP and mCherry filters (Nippon Roper, Tokyo, Japan), a CSU22 scanning system (Yokogawa Electronics, Tokyo, Japan), and an Andor iXon cooled digital CCD camera (Andor Technology PLC, Belfast, UK). The captured images were analyzed using Andor iQ software version1.9 (Andor Technology PLC, Belfast, UK).

### ROS accumulation detection

The intracellular levels of ROS were detected using the oxidant-sensing probe, 2’,7’-dichlorodihydrofluorescein diacetate (H_2_DCFDA, Molecular Probes, Eugene, OR, USA). Conidia were incubated in an optimized minimal medium for 20 h at 25°C. The culture was washed with distilled water and incubated with 0.01 mM H_2_DCFDA for 30 min at 25°C. This was followed by washing the culture twice with distilled water, suspension in 20 mL MOPS7 buffer (0.1 M MOPS, pH 7.0), and observation by fluorescence microscopy.

## ACKNOWLEDGMENTS

This study was financially supported by Japan Society for the Promotion of Science (JSPS) KAKENHI Grant Number JP17K15242, Grant-in-Aid for Young Scientists (B), and 18H02123, Grant-in-Aid for Scientific Research (B).

## Figure Legends

**FIG S1. Phylogenetic tree of *A. oryzae* strains**. Phylogenetic tree of *A. oryzae* strains classified based on comparative genomic hybridization analysis (http://nribf21.nrib.go.jp/CFGD/). All the strains are classified into 13 phylogenetic clades based on the phylogenetic distances. The strains used in this study are presented in bold letters.

**FIG S2. Correlation between heterokaryon incompatibility and strain phylogeny in *A. oryzae***. (A) VCG classification method based on auxotrophic complementation assay. (B) Analysis of incompatibility among 13 inter-clade strains. The 13 representative strains belong to 13 different clades. Protoplasts in self- or non-self-pairing with different uridine/uracil auxotrophies (Δ*pyrF* and Δ*pyrG*) were forcedly fused using polyethylene glycol (PEG). The fused protoplasts were plated on the CD minimal agar medium without uridine/uracil at 30°C for 5 days. Self-pairing, which served as the positive control, is shown at the diagonal line. Only the non-self-pairing of RIBOIS01-RIB143 and RIB40-RIB81 generated auxotrophically complemented heterokaryons. The rest did not form any significant hyphae. All the pairings were performed at least in triplicate. (C) Incompatibility analysis of intra-clade strains. Strains belonging to the same clade were selected, e.g., two strains in clade 1, 11 strains in clade 3, five stains in clade 4, three strains in clade 5, three strains in clade 8, four strains in clades 9 and 10 (owing to the compatibility between the RIB40 strain of clade 10 and the RIB81 strain of clade 9).

**FIG S**3. **ROS accumulation and fluorescence visualization of physiological responses in self and non-self-pairings**. (A) Construction of fluorescence strains expressing EGFP and mCherry in the cytoplasm. (B) Gradual cytoplasmic mixing of EGFP and mCherry in heterokaryotic cells following the self-pairing of RIB640-Δ*pyrF*-EGFP and RIB640-Δ*pyrG*-mCherry strains. Arrowheads and white arrows indicate fusion points and septa, respectively. (C) ROS accumulation in the fused area of incompatible pairings between RIB640Δ*pyrF* and RIB40Δ*pyrG*-mCherry, as well as RIB640Δ*pyrF* and RIB640Δ*pyrG*-mCherry. (D) Construction of fluorescence strains expressing EGFP and mCherry in the nuclei, ER, and mitochondria. Scale bars represent 10 μm.

**FIG S4. Visualization of physiological responses in nuclei**. (A) Self pairing between RIB40Δ*pyrF*-EGFP and RIB40Δ*pyrG*-H1-mCherry. (B) Self pairing betweenRIB640Δ*pyrF*-EGFP and RIB640Δ*pyrG*-H1-mCherry. (C) Incompatible pairing between RIB40Δ*pyrF*-EGFP and RIB640Δ*pyrG*-H1-mCherry. Scale bars represent 10 μm.

**FIG S5. Visualization of physiological responses in the endoplasmic reticulum**. (A) Self pairing between RIB40Δ*pyrF*-AoBipA-EGFP and RIB40Δ*pyrG*-mCherry. (B) Self pairing between RIB640Δ*pyrF*-AoBipA-EGFP and RIB640Δ*pyrG*-mCherry. (C) Incompatible pairing between RIB40Δ*pyrF*-AoBipA-EGFP and RIB640Δ*pyrG*-mCherry. Note that incompatible fusion was started after the 0-min time point as indicated by the entry of mCherry fluorescence into the cells of the AoBipA-EGFP-labelled strain. Scale bars represent 10 μm.

**FIG S6. Visualization of physiological responses in mitochondria following incompatible pairing**. (A) Self pairing between RIB640-Δ*pyrF*-AoCit1-EGFP and RIB640-Δ*pyrG*-mCherry. (B) Incompatible pairing between RIB640Δ*pyrF*-AoCit1-EGFP and RIB40Δ*pyrG*-mCherry.

**FIG S7. Alleviation of incompatibility-triggered cell death by deleting mitochondrial fission-related genes**. (A) Construction of *Aofis1*/*Aodnm1* deletion strains using CRISPR/Cas9 system. The DNA electrophoresis and growth phenotype were examined in each strain with *Aofis1*/*Aodnm1* deletions. (B) Incompatible pairing with *Aofis1*/*Aodnm1* deletions was performed via the protoplast fusion assay to check whether incompatibility was blocked or not. (C) PI staining of grown heterokaryons derived from forcedly fused self/incompatible protoplasts with *Aofis1*/*Aodnm1* deletions. Scale bars represent 10 μm.

**FIG S8**. (A) Construction of RIB40Δ*pyrF*Δ*Aofis1*-EGFP, RIB40Δ*pyrF*Δ*Aodnm1*-EGFP, RIB40Δ*pyrG*Δ*Aofis1*-mCherry, RIB640Δ*pyrG*Δ*Aofis1*-mCherry, RIB40Δ*pyrG*Δ*Aodnm1*-mCherry, and RIB640Δ*pyrG*Δ*Aodnm1*-mCherry. (B) Construction of RIB40Δ*pyrF*Δ*Aofis1*-AoCit1-EGFP and RIB40Δ*pyrF*Δ*Aodnm1*-AoCit1-EGFP. Scale bars represent 10 μm.

